# Eye and hand movements disrupt attentional control

**DOI:** 10.1101/2020.09.22.309237

**Authors:** Nina M. Hanning, Luca Wollenberg, Donatas Jonikaitis, Heiner Deubel

## Abstract

Voluntary attentional control is the ability to selectively focus on a subset of visual information in the presence of other competing stimuli. While it is well established that this capability is a marker of cognitive control that allows for flexible, goal-driven behavior, it is still an open question how robust it is. In this study we contrasted voluntary attentional control with the most frequent source of automatic, involuntary attentional orienting in daily life—shifts of attention prior to goal-directed eye and hand movements. In a multi-tasking paradigm, we asked participants to attend at a location while planning eye or hand movements elsewhere. We observed that voluntary attentional control suffered with every simultaneous action plan. Crucially, this impairment occurred even when we reduced task difficulty and memory load—factors known to interfere with attentional control. Furthermore, the performance cost was limited to voluntary attention. We observed simultaneous attention benefits at two movement targets without attentional competition between them. This demonstrates that the visual system allows for the concurrent representation of multiple attentional foci. It further reveals that voluntary attentional control is extremely fragile and dominated by automatic, premotor shifts of attention. We propose that action-driven selection disrupts voluntary attention and plays a superordinate role for visual selection.

## Introduction

Attentional control is the ability to select relevant visual information in the presence of other, non-relevant stimuli [Posner, 1980; Treisman, 1982]. This selection is also referred to as top-down, or task-driven attention, and can be contrasted to bottom-up attention which automatically selects stimuli based on their unique properties [Müller & Rabbitt, 1989; Theeuwes, Kramer, Hahn, Irwin, 1998; Klein, 2000; Carrasco, 2004; Carrasco, 2011]. Top-down selection is typically investigated by having humans and non-human animals attend to one out of several stimuli, either by instruction or manipulating reward probabilities [Found & Müller, 1996; Ciaramitaro, Cameron, Glimcher, 2001; Anderson, Laurent, Yantis, 2011; Baruni, Lau, Salzman, 2015]. Generally, attentional control is proposed to play a crucial role in the transition from automatic to flexible, adaptive behavior [Aston-Jones, Desimone, Driver, Luck, & Posner, 1999; Munoz & Everling, 2004; Schütz, Trommershäuser, & Gegenfurtner, 2012].

A separate line of research has focused on visual attention in the context of motor actions. Eye movements [Deubel & Schneider, 1996; Hoffman & Subramaniam, 1995; Kowler, Anderson, Dosher, & Blaser, 1995; Montagnini & Castet, 2007; Jonikaitis, Klapetek, Deubel, 2017; Hanning, Szinte, Deubel, 2019] as well as hand movements [Deubel, Schneider, & Paprotta, 1998; Baldauf, Wolf, & Deubel, 2006; Hesse & Deubel 2011; Rolfs, Lawrence & Carrasco, 2013] are preceded by shifts of attention to their motor targets. These premotor shifts of attention are thought to occur automatically when we explore or interact with our environment; in other words, without any instructions or reward manipulations [Baldauf & Deubel, 2010]. Attentional control and premotor attention frequently share a common goal: Behaviorally relevant or rewarded objects are typically also motor targets [Land & Lee, 1994; Land & McLeod, 2000]. However, given that premotor attention is thought to be automatic while attentional control is task-driven and voluntary, the relationship between the two has not yet been defined.

Several frameworks can be used to describe the relationship between attentional control and premotor attention. The key approach to study attentional control is the *visual search* paradigm, which requires top-down or voluntary attention to select task relevant information [Treisman, 1982]. Unfortunately, visual search paradigms frequently require gaze to be maintained stable and do not measure eye movements. They typically assume that, under free-viewing conditions, gaze would be directed to the attended location [McPeek, Skavenski, Nakayama, 2000; Peters, Iyer, Itti, & Koch, 2005; Bichot, Rossi, & Desimone, 2005]. Thus, even though it is implied in these studies that attentional control drives premotor attention, the relationship between the two is not directly investigated. Based on the visual search paradigm, a more formal framework has been proposed, referred to as *salience maps or priority maps* [Itti & Koch, 2001; Thompson & Bichot, 2005; Fecteau & Munoz, 2006; Bisley & Goldberg, 2010]. Within these maps, bottom-up and top-down signals are thought to be integrated in a winner-take-all process. Subsequently, the highest activity peak on the map determines the attentional focus, to which eye movements can be potentially directed [Itti & Koch, 2001]. This framework again links attentional control and premotor attention without explicitly testing their relationship.

While visual-search-based theories assume that eye movements *follow* the attentional focus, the *premotor theory of attention* proposes the opposite, namely that visual attention is a product of the motor system. In order to shift attention covertly (i.e., without moving the eyes) a motor program still has to be prepared, yet not necessarily executed [Rizzolatti, Riggio, Dascola, Umiltá, 1987; Craighero, Fadiga, Rizzolatti, Umiltà, 1999]. While this theory can explain automatic attention shifts to motor targets in the absence of any instruction or reward manipulation, a tight coupling does not prove that visual attention in fact arises from motor preparation. Instead, the reverse might be true, and visual attention is necessary to successfully perform a motor action. This alternative account states that attentional selection is required to specify the motor coordinates for an upcoming movement [Schneider, 1995; Baldauf & Deubel, 2010]. Baldauf and Deubel (2010) proposed an *attentional landscapes* framework which explicitly deals with multiple attentional foci, as they can occur during simultaneous eye-hand movements [Jonikaitis & Deubel, 2011; Hanning, Aagten-Murphy, & Deubel, 2018; Kreyenmeier, Deubel, & Hanning, 2020]. While cognitive control and premotor attention are closely linked in both frameworks, the implied direction of this relationship is opposite.

The above discussed frameworks vary markedly in their assumptions. The relationship between attentional control and premotor attention is either not directly specified (visual search & priority maps), or attention is necessarily driven by motor selection (premotor theory of attention), or, the contrary, motor selection requires attention, but not the other way around (attentional landscapes). Crucially, these contrasting assumptions have not yet been addressed. It is therefore still an open question how reflexive, automatic information selection in the context of motor actions interacts with adaptive, controlled attentional selection.

We investigated these two components of attentional selection using a classical dissociation approach in which we pitted premotor attention shifts against the capability to maintain voluntary spatial attention. This approach can reveal competition or prioritization between premotor and voluntary attention. Our participants were required to attend to a given location (*voluntary attention*) while simultaneously preparing an eye movement (*premotor attention to eye*) and/or hand movement (*premotor attention to hand*) to another location. If all three tasks interact equally with each other, this would indicate dual-task costs, whereas distinct interaction patterns can differentially support or refute the above discussed frameworks. We used local visual discrimination performance as a proxy of visuospatial attention during premotor and voluntary selection, and systematically biased participants’ deployment of voluntary attention by informing them about which location was most likely to contain the discrimination signal—a briefly presented oriented noise patch. Our data revealed that any type of attentional selection, voluntarily as well as premotor, was associated with improved discrimination performance at the target location. Furthermore, we observed no indication of attentional competition between the eye and the hand motor target. In striking contrast, voluntary attentional selection suffered with every motor action being planned, revealing that eye and hand movement preparation abolishes attentional control. This demonstrates that the visual system selectively prioritizes automatic shifts of attention to motor targets over top-down attentional control.

## Results

In **Experiment 1** participants were instructed to perform different combinations of three possible tasks: endogenously attending to a specific location (*Attention*), executing an eye movement (*Eye*), and executing a hand movement (*Hand*) to a centrally cued target. Concurrently, they performed a two-alternative forced-choice discrimination task (***Figure 1a***) based on oriented pink noise patches [Hanning, Deubel, Szinte, 2019] (***Figure 1b***). Orientation discrimination performance at the endogenously attended location, at the motor target location(s), and at neutral locations (i.e., movement-irrelevant, non-target control locations) served as a proxy for visuospatial attention during motor target and endogenous perceptual selection. Altogether, the experiment comprised seven tasks: *Attention-only, Eye-only, Hand-only, Eye-Hand, Attention-Eye, Attention-Hand, Attention-Eye-Hand* (see ***Figure 1c*** for stimulus timing).

**Figure 1.**
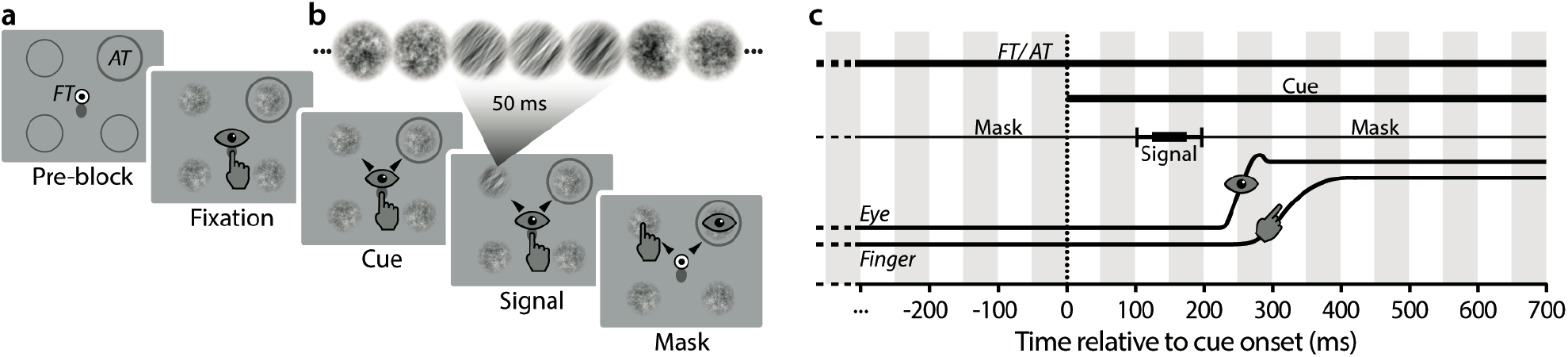
Methods. (**a**) Example trial sequence of the *ATT-EYE-HAND* task (Experiment 1). Throughout the block, the endogenous attention target (AT) was marked by a dark gray circle. Participants maintained central eye and finger fixation until two black arrow cues occurred that marked two of four noise patches as motor targets. Participants reached towards one and simultaneously saccaded towards the other motor target. Before movement onset, one of the noise streams showed a clockwise or counterclockwise orientation signal. After the movements and a masking period, participants indicated their discrimination judgement via button press. (**b**) Noise streams used as discrimination stimuli. Each of the four noise streams consisted of a succession of randomly generated 1/f noise patches.The test stream comprised a 50 ms sequence of orientation filtered 1/f noise patches showing a clockwise or counterclockwise tilt. (**c**) Stimulus timing. Fixation (FT) and attention target (AT) remained on the screen throughout the trial. 400 to 800 ms after the onset of four noise pre-masks (M), the motor cues were presented. 100 ms after cue onset, one of the noise streams contained the orientation test signal, which was masked after 50 ms.

First, we measured the pattern of attentional selection during each of the above conditions (***Figure 2a***). In the *Attention-only* task, we biased discrimination signal probability to guide voluntary attention: the discrimination signal was most likely to appear at the to be attended location (75% probability). Performance at the attention target was better than at the non-targets (*p =* 0.001), indicating that participants deployed voluntary attention to the most probable discrimination signal location [Carrasco, 2004; Müller & Rabbit, 1989]. In the *Eye-only* and the *Hand-only* task, performance at the eye target (*p =* 0.001) and the hand target (*p* = 0.004) was similarly enhanced relative to the movement-irrelevant locations which were equally likely to contain the discrimination signal. This demonstrates that attention shifted automatically to the motor targets, independent of discrimination signal probability.

**Figure 2.**
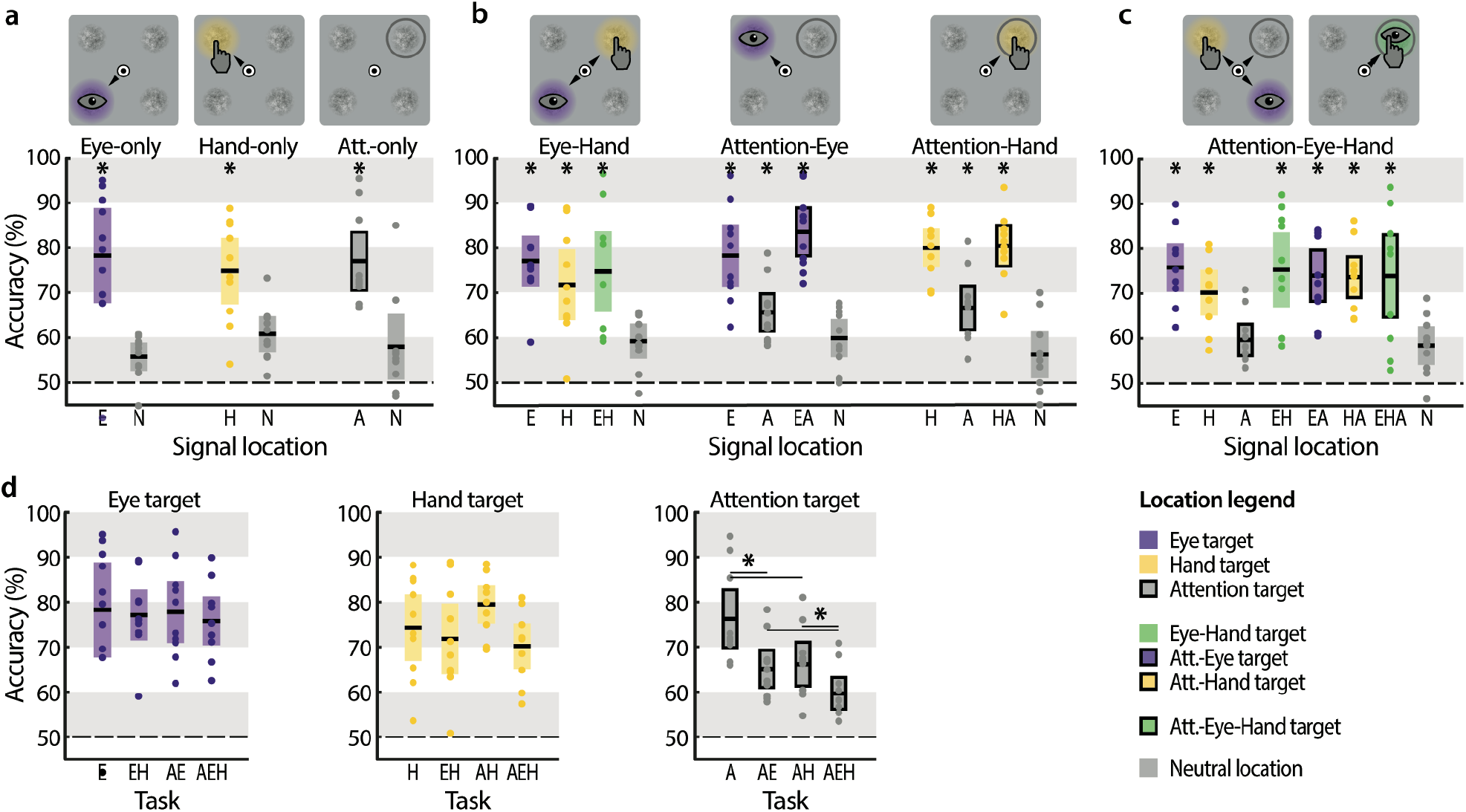
Voluntary and premotor attentional selection. Discrimination performance at the different test locations of the single target tasks (**a**), double target tasks (**b**), and tripe target task (**c**) of Experiment 1. Black lines within each whisker plot indicate the group average. Colored bars depict the 95% confidence interval, dots represent individual subject data, and dashed lines mark chance performance. \**p* < 0.05, significant difference between a location and the neutral location of the respective task. (**d**) Discrimination performance at the eye target (left), hand target (middle), and attention target (right) as a function of the experimental task. \**p* < 0.05, all significant difference between tasks are indicated by horizontal lines. Other conventions as in (a).

Next, we investigated interactions between these three sources of attentional selection (***Figure 2b***). In the *Eye-Hand* task, participants simultaneously performed eye and hand movements to either shared or separate targets. When the two movements were made to separate target locations, we observed improved performance both at the eye (*p =* 0.003) and the hand target (*p* = 0.007) compared to the movement-irrelevant locations, and the attentional benefit at the two motor targets did not differ (*p* = 0.219). When participants made simultaneous eye-hand movements to a shared target, performance at that location was also significantly improved (*p* = 0.005), and comparable to performance when eye and hand movements were directed to separate locations (compared to the eye target: *p* = 0.431; compared to the hand target: *p* = 0.515). Importantly, relative to single effector movements (i.e. *Eye-only* and *Hand-only*), combined effector movements to separate locations did neither reduce discrimination performance at the eye target (*p* = 0.819) nor at the hand target (*p* = 0.366). In summary, this demonstrates that during simultaneous eye-hand movements, attention is deployed to both motor targets in parallel without any observable cost, which is in line with previous studies [Jonikaitis & Deubel, 2011; Hanning et al., 2018; Kreyenmeieret al., 2020; but see Khan, Song, & McPeek, 2011].

To investigate how voluntary attentional control interacts with motor planning, we asked participants to attend at one location while preparing an eye or hand movement to another (*Attention-Eye* task, *Attention-Hand* task). These two tasks create a conflict: while the discrimination signal was most likely to appear at the voluntary attention target, the movement target was more likely to be at a different, non-predictable location. In the *Attention-Eye* task, when eye movement target location and voluntarily attended location coincided, this mutual target, as expected, received a discrimination benefit (*p* = 0.001). When voluntary attention and eye movement were directed to separate locations, we observed enhanced performance at the eye target (*p* = 0.001) and a small performance benefit at the attention target (*p* = 0.015; note that *p*_*corrected*_ = 0.060). Moreover, performance at the attention target was worse than at the eye target (*p* = 0.001). We observed similar results for voluntary attention during hand movement preparation. In the *Attention-Hand* task, performance at the attention target (*p* = 0.007), the hand target (*p* = 0.001), and the shared hand-attention target (*p* = 0.001) was significantly enhanced. Again, the attentional benefit at the attention target was smaller than at the hand target (*p* = 0.001). To summarize, in contrast to the *Eye-Hand* task, in which attention was equally distributed to both motor targets, attention was clearly biased towards the motor target in the *Attention-Eye* and the *Attention-Hand* tasks.

We put further stress on attentional control by asking participants to simultaneous attend to a location while preparing both an eye and a hand movement (***Figure 2c***, *Attention-Eye-Hand*). As before, we observed a clear attentional benefit at the eye target (*p* = 0.001), the hand target (*p* = 0.002), and the combined eye-hand target (*p* = 0.001). However, even though the discrimination signal was most likely to appear at the voluntary attention target, participants were not able to maintain voluntary attention there (*p* = 0.534).

A direct comparison of performance across the different motor tasks showed that this decrease in performance was limited to voluntary attention and did not apply to motor targets (***Figure 2d***). Performance at the eye target was consistently enhanced whether only an eye movement was prepared, or the eye movement was accompanied by either a hand movement (*p* = 0.819), voluntary attentional selection (*p* = 0.893), or both (*p* = 0.645). Likewise, attention was consistently deployed to the hand target, independently of whether the hand movement was accompanied by an eye movement (*p* = 0.366), voluntary attentional selection (*p* = 0.180), or both (*p* = 0.150). In other words, performance at the motor targets in the combined eye-hand movement task was indistinguishable from the respective performance in the single (eye only or hand only) tasks, demonstrating that the attentional selection of one motor target did not affect the selection of the other. In direct contrast, voluntary attentional control was hampered by motor programming: performance at the attention target was reduced whenever a single eye movement (*p* = 0.001) or single hand movement (*p* = 0.003) were planned. Importantly, performance decreased even further when both an eye and a hand movement simultaneously were directed away from the attended location (compared to single eye movement: *p* = 0.001; compared to single hand movement: *p* = 0.009). Thus, while attentional control was already affected by single movements, it was practically annihilated during simultaneous eye and hand movement preparation.

We observed that voluntary attention was reduced when participants made an eye or hand movement. Conversely, however, voluntary attention did not affect perceptual performance at the motor targets. We next investigated whether voluntary attention interfered with eye or hand movement preparation in any other way, for example by decreasing movement accuracy or prolonging movement latencies.

We first compared eye and hand landing positions across the different motor tasks (***Figure 3a***). Generally, when two motor targets were cued (*Eye-Hand* and *Attention-Eye-Hand* task) participants tended to select the upper locations as eye targets and the lower locations as hand targets. However, neither eye nor hand movement precision—measured as the average distance of the movement endpoint from motor target center—differed between the respective single movement tasks (*Eye-only* / *Hand-only*) and the multiple target tasks (Eye movement precision: *Eye-only* vs. *Eye-Hand*: *p* = 0.229*3, vs. *Attention-Eye*: *p* = 0.957*3, vs. *Attention-Eye-Hand*: *p* = 0.308*3; Hand movement precision: *Hand-only* vs. *Eye-Hand*: *p* = 0.060*3, vs. *Attention-Hand*: *p* = 0.721, vs. *Attention-Eye-Hand*: *p* = 0.037). Thus, neither the requirement to program a second movement nor to deploy voluntary attention affected eye and hand movement precision. In contrast, we observed interactions between eye and hand movement control with respect to movement latencies. Compared to the *Eye-only* task (***Figure 3b***; left), eye movement onsets were significantly delayed in tasks in which also a hand movement had to be prepared (*Eye-only* vs. *Eye-Hand*: *p* = 0.001, vs. *Attention-Eye-Hand*: *p* = 0.001), which is in line with earlier work [Jonikaitis & Deubel, 2011]. Having to attend voluntarily, however, did not slow down eye movement execution (*Eye-only* vs. *Attention-Eye*: *p* = 0.106). Likewise, hand movement latencies (***Figure 3b***; right) were slightly prolonged by simultaneous eye movement preparation (*Hand-only* vs. *Eye-Hand*: *p* = 0.019, vs. *Attention-Eye-Hand*: *p* = 0.012), but again not by voluntary attention (*Hand-only* vs. *Attention-Hand*: *p* = 0.301). Note that the effect of hand movement on eye movement execution was considerably more pronounced than vice versa.

**Figure 3.**
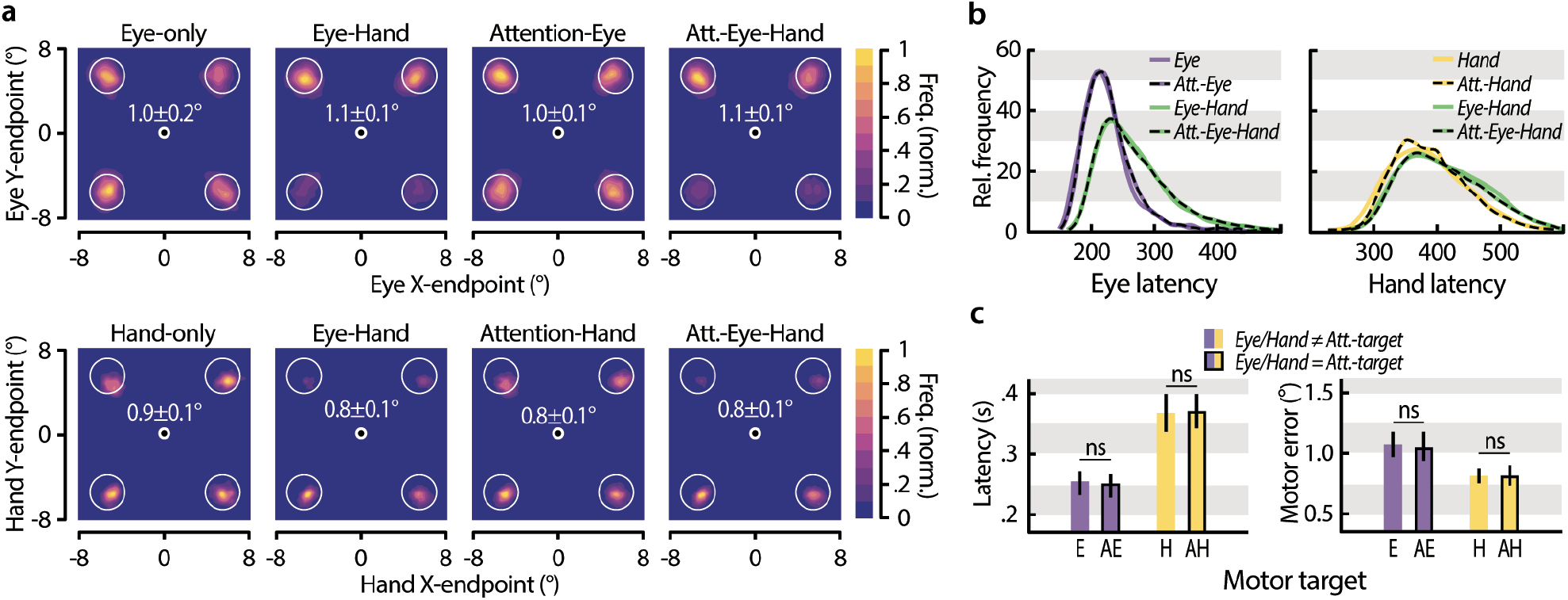
Movement latencies and precision. (**a**) Normalized eye (top row) and hand (bottom row) landing frequency maps averaged across participants in the respective tasks. White values depict the average distance between movement endpoint and target center as well as the the 95% confidence interval. (**b**) Relative frequency of eye (left) and hand right) latencies in the respective tasks. (**c**) Eye and hand movement latencies (left) landing errors (middle) and endpoint variance (right) across all tasks split as to whether the movement was made to an endogenously attended location (AE, AH) or not (E, H). Error bars denote the 95% confidence interval.

To investigate the influence of voluntary attentional control on movement execution, we evaluated movement latencies (***Figure 3c***; left) and landing errors (defined as the distance between movement endpoint and target center; ***Figure 3c***; right) depending on whether the movement was made to the voluntarily attended location or not. Neither for eye nor for hand movements we observed a significant difference in latencies (*Eye-only* vs. *Attention-Eye*: *p* = 0.164*1; *Hand-only* vs. *Attention-Hand*: *p* = 0.118*1) or landing errors (*Eye-only* vs. *Attention-Eye*: *p* = 0.517; *Hand-only* vs. *Attention-Hand*: *p* = 0.865*1), demonstrating that attentional control affected neither movement onset nor precision.

Our results showed that preparing eye or hand movements interferes with voluntary attention. In an attempt to reduce the interference of motor preparation on attentional control, we optimized conditions to favor voluntary attention deployment in **Experiment 2**. In this experiment, we presented only one noise stream that always contained the discrimination signal, which removes any potential uncertainty as to where to attend or respond (***Figure 4a***). Participants either attended to that location (*Attention-only*), or attended and made eye (*Attention-Eye*), hand (*Attention-Hand*), or simultaneous eye-hand movements (*Attention-Eye-Hand*) away from this location. In this experiment, we varied the width of the orientation filter used to create the discrimination signal (the smaller the width, the clearer the orientation) and assessed perceptual performance by measuring psychometric thresholds—an alternative approach to quantify attention [Carrasco, 2011].

**Figure 4.**
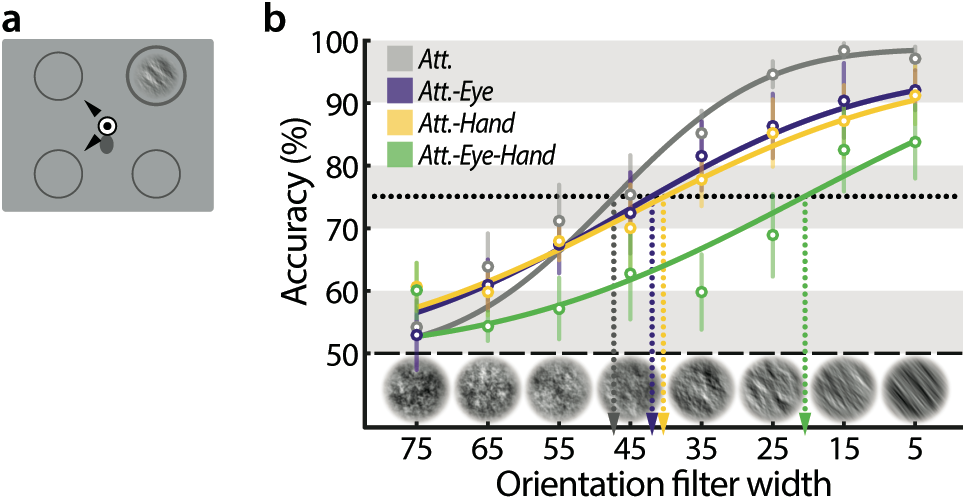
Costs of action-selection for voluntary attention. (**a**) Stimuli configuration in Experiment 2. One noise stream was presented at a fixed location, marked by a circle. Participants maintained central eye and finger fixation until one (*Attention-Eye* or *Attention-Hand*) or two (*Attention-Eye-Hand*) black arrow cues occurred, indicating the motor target(s). Participants reached and / or saccaded towards the motor target(s) upon cue onset. In the *Attention* task no cue occurred and participants maintained fixation. After 100 −200 ms the noise stream displayed a brief orientation, which participants discriminated via button press at the end of the trial. (**b**) Group-averaged psychometric functions (discrimination performance vs. orientation filter width) for each task. Error bars denote the between subject SEM.

In line with our previous findings, motor preparation consistently reduced participants’ ability to voluntary attend (***Figure 4b***). This was evident in the psychometric functions’ slopes and thresholds. In the *Attention-only* task, the slope (*m* = 0.59 [0.37, 1.58]) was steeper than in the *Attention-Eye* task (*m* = 0.37 [0.26, 0.65]) and the *Attention-Hand* task (*m* = 0.32 [0.17, 0.57]). In the hardest version of the task (*Attention-Eye-Hand*), the detrimental effect of motor-preparation on attentional control was even more pronounced (*m* = 0.29 [0.07, 1.38]). When estimating thresholds at a fixed discrimination performance level of 75% we observed a matching pattern: the highest threshold in the *Attention-only* task (*α* = 47.21 [41.89, 53.51]), followed by the two single movement tasks (*Attention-Hand*: *α* = 42.09 [31.11, 49.12], *Attention-Hand*: *α* = 40.16 [31.40, 47.22]). Correspondingly, performance in the *Attention-Eye-Hand* task yielded the lowest threshold (*Attention-Eye-Hand*: *α* = 20.50 [0.98, 36.44]). In summary, even though we provided optimal conditions for voluntary attentional control by decreasing task difficulty and memory load, motor preparation nonetheless markedly impaired voluntary attentional selection.

## Discussion

We studied the relationship between voluntary attentional selection and automatic attention shifts before goal-directed motor actions. We observed robust perceptual benefits (a marker for attention deployment) under the typically investigated single-task conditions: at eye movement targets [Deubel & Schneider, 1996; Hoffman & Subramaniam, 1995; Kowler et al., 1995; Montagnini & Castet, 2007; Jonikaitis, Klapetek, Deubel, 2017; Hanning, Szinte, Deubel, 2019], hand movement targets [Deubel et al., 1998; Baldauf et al., 2006; Rolfs et al., 2013], as well as endogenously attended locations [Posner, 1980; Müller & Rabbitt, 1989; Montagna, Pestilli, & Carrasco, 2009]. Under multiple-task conditions, when participants either had to select two or even three locations, we observed a dissociation between the investigated sources of attention. There was no attentional competition between the eye and the hand motor targets. In striking contrast, voluntary attentional selection suffered from every simultaneous action plan.

Our results reveal that the visual system selectively prioritizes automatic, premotor shifts of attention over voluntary attentional control, and demonstrate that voluntary attentional selection is extremely fragile. Even though task instructions and stimulus probabilities particularly facilitated voluntary attentional orienting, it was consistently susceptible to disruption by both eye and hand movements. Indeed, one could have expected the opposite, namely that in order to maintain attentional control, discrimination performance should have dropped at motor goals.

Earlier studies observing that eye movements compete with voluntary attention or vice versa [Montagnini & Castet, 2007; Deubel, 2008] cannot rule out that participants prioritized one task over the other due to increased difficulty (dual-as compared to a single-task condition). Using different levels of task difficulty (one to three attention targets), we show that voluntary attention was increasingly impaired, whereas, crucially, performance at eye and hand movement targets did not suffer. This dissociation—a cost for endogenous attention yet no interference for eye and hand premotor attention—rules out that increased task or attentional selection difficulty can explain the loss of attentional control. Further, even when there was only a single location to be voluntarily attended (Experiment 2), we observed a decrease in attentional control whenever an eye or hand movement was planned away from this location. This shows that neither reduced stimulus location uncertainty nor the attention target being the only salient stimulus presented could prevent the observed loss of attentional control.

Different aspects of our results are not compatible with other major frameworks referring to the relationship between attentional control and premotor attention. First, our and earlier observations of multiple, simultaneous attentional peaks [Baldauf & Deubel, 2009; Jonikaitis & Deubel, 2011; Gilster, Hesse, & Deubel, 2012; Hanning et al., 2018; Kreyenmeier et al., 2020], are not compatible with *priority map* models assuming a strict winner-take-all attentional selection, in which performance benefits should occur only at the highest peak [Itti & Koch, 2001; Thompson & Bichot, 2005; Fecteau & Munoz, 2006; Bisley & Goldberg, 2010]. Second, our finding that eye and hand movements draw attention away from the voluntary attended location is incompatible with the view that motor actions merely follow the current focus of voluntary attention, as is assumed in visual search frameworks [Treisman, 1982; Itti & Koch, 2001; Müller & Krummenacher, 2006; Theeuwes & Burger, 1998]. Third, the *premotor theory of attention* [Rizzolatti et al., 1987; Craighero et al., 1999] assumes that any shift of attention is equivalent to a saccade plan, which offers two testable predictions. First, when the saccade target matches the endogenously attended location, saccadic latencies should be shorter. Our data show that they are not. Second, when the saccade is directed away from the endogenously attended location, two “saccade plans” are technically required. As these should compete with each other, attention allocation to both locations should decrease. Our data show that this also was not the case. In summary, our findings cannot be explained by priority map, visual search, or the premotor theory of attention frameworks without modifying their core assumptions.

Our findings are in line with the proposal of an *attentional landscape*—a map representing the attentional distribution across space [Baldauf & Deubel, 2010]. This framework allows for a simultaneous deployment of visual attention to several action-relevant locations, observed as multiple “attentional peaks”. These peaks can vary in magnitude to reflect different degrees of attentional allocation. Indeed, our results demonstrate that attention can be allocated to multiple locations at a time. Multiple stimuli are typically assumed to compete for attentional selection [Bundesen, 1990; Bundesen & Habekost, 2014]. We found that motor goals are prioritized in this attentional selection. Further in line with this view, we observed that the peak performance associated with endogenous attentional selection decreases gradually with every motor action added.

Some of our behavioral effects can be linked to earlier neurophysiological studies. First, activity within visual areas is consistently modulated by visual attention [Moran & Desimone, 1985; McAdams, Maunsell, 1999], and this modulation is thought to result in corresponding changes in perceptual performance [Moore & Fallah, 2004; Lovejoy & Krauzlis, 2010; Fernández & Carrasco, 2020]. Several studies focusing on voluntary attention, or attentional control, have observed this modulation throughout the hierarchy of the visual system, ranging from occipital [Mazer & Gallant, 2003; Reynolds & Chelazzi, 2004; Serences, 2008] over parietal [Colby, Duhamel, & Goldberg, 1996; Ipata, Gee, Bisley, & Goldberg, 2009] to frontal cortex [Juan, Shorter-Jacobi, & Schall, 2004; Schafer & Moore, 2007; Cosman, Lowe, Zinke, Woodman, & Schall, 2018; Jonikaitis & Moore, 2019]. Second, eye movement preparation modulates neuronal activity in visual cortical areas in a manner indistinguishable from voluntary attention [Moore & Armstrong, 2003]. The neural sources for this modulation are assumed to be fronto-parietal feedback connections converging onto earlier visual areas [Moore & Fallah, 2004; Armstrong & Moore, 2007; Ekstrom, Roelfsema, Arsenault, Bonmassar, & Vanduffel, 2008; Rolfs & Szinte, 2016]. This has led to multiple proposals that oculomotor areas could serve as an attentional source or map [Fecteau & Munoz, 2006; Bisley & Goldberg, 2010; Bisley & Mirpour, 2019]. Third, our behavioral observation of multiple attentional peaks can be related to simultaneous and distinct activity peaks observed in human and monkey neurophysiology studies [Morawetz, Holz, Baudewig, Treue, & Dechent, 2007; Baldauf, Cui, & Andersen, 2008; Niebergall, Khayat, Treue, & Martinez-Trujillo, 2011; Steinmetz & Moore, 2014; Saber, Pestilli, & Curtis, 2015].

The neurophysiological basis for other key aspects of our findings however is still lacking. First, there is no evidence of the source of premotor attention signals to visual cortex before hand movements. These feedback-signals could originate from reach-related or oculomotor areas—but this has not yet been investigated. Thus, it is not known whether neuronal activity associated with premotor attention before eye and hand movements occurs simultaneously in the same area, or in separate areas. Given how consistently attention is shifted before reaching [Deubel et al., 1998; Baldauf et al., 2006; Rolfs et al., 2013] and grasping movements [Hesse & Deubel, 2011; Hesse, Schenk, & Deubel, 2012], understanding the underlying neural circuitry is crucial to comprehend the mechanisms that govern attentional selection in real-life situations. Second, interactions between endogenous and premotor attention likewise have not yet been explained neurophysiologically. Oculomotor areas are thought to be the common source for presaccadic and covert voluntary attention [Moore & Fallah, 2004; Armstrong & Moore, 2007; Ekstrom et al., 2008]. It is therefore unclear why both eye movement and hand movement planning do compete with voluntary attention, but no competition is observed between multiple motor targets [Godijn, Theeuwes, 2003; Baldauf & Deubel, 2008; Baldauf & Deubel, 2009; Jonikaitis & Deubel, 2011; Hanning et al., 2018; Kreyenmeier et al., 2020]. Third, we do not know whether the neuronal modulations associated with eye, hand, and voluntary spatial attention can be observed in a common area (e.g., frontal, parietal, visual or subcortical areas), suggesting a common attentional map, or whether they arise from different areas. In the latter case, attention to multiple targets could activate separate areas without integrating activity between them (and thus limiting premotor competition between attentional goals). As evidence in favor of a common attentional map has mainly been collected under experimental conditions requiring eye and hand fixation, those conclusions may be biased. It is equally possible that separate, effector-specific maps show attentional modulation during eye and hand movement target selection. Such separate maps could explain the absence of premotor attentional competition between different effectors. In sum, our results show that there are multiple questions to be answered if one wants to explain everyday attentional selection.

In everyday life we continuously explore and interact with our environment. Our findings reveal that whenever our eye or hand movement goals do not match our attentional control settings, attentional control cannot be maintained. Thus, attentional control is likely to fail as frequently as we move. We typically avoid this failure by aligning our attentional control and movement goals. While the classical understanding of attention underscores covert attentional orienting in the absence of motor actions, such situations of immobility are rare, if not artificial. Actions are typically considered the *consequence* of attentional control. Our data however show that actions take precedence over attentional control. We therefore propose to refocus from considering action as the strict consequence of voluntary attentional control to viewing action as the main determinant of successful or failed visual selection.

## Materials and Methods

### Participants and setup

The sample sizes were determined based on previous work [Jonikaitis & Deubel, 2011; Hanning et al., 2018]. Ten participants (ages 23–31 years, 7 female) completed Experiment 1, six participants (ages 23–28 years, 4 female) took part in Experiment 2. All participants were healthy, had normal vision and, except for one author (N.M.H.), were naive as to the purpose of the experiments.

Gaze position was recorded using an EyeLink 1000 Tower Mount (SR Research, Osgoode, Ontario, Canada) at a sampling rate of 1 kHz. Manual responses were recorded via a standard keyboard. The experimental software controlling display, response collection, as well as eye tracking was implemented in Matlab (MathWorks, Natick, MA, USA), using the Psychophysics [Brainard, 1997; Pelli, 1997] and EyeLink toolboxes [Cornelissen, Peters, & Palmer, 2002]. Stimuli were presented on a 45° inclined touchscreen (Elo 2700 IntelliTouch, Elo Touchsystems, Menlo Park, CA) with a spatial resolution of 1280×1024 pixels and a refresh rate of 60 Hz.

### Experimental Design

**Experiment 1** comprised seven tasks (randomized block design): *Attention-only, Eye-only, Hand-only, Eye-Hand, Attention-Eye, Attention-Hand, Attention-Eye-Hand*. ***Figure 1a*** depicts the sequence for the *Attention-Eye-Hand* task: Participants initially fixated a central fixation target (FT) comprising a black (∼0 cd/m2) and white (∼98 cd/m2) “bull’s eye” (radius 0.5°) on a gray background (∼60 cd/m2). Their right index finger remained on a gray oval (0.6° x 0.65°, ∼22 cd/m2) slightly below the eye fixation. At the beginning of each block, four equally spaced locations were marked by gray circles (radius 1.7°) at an eccentricity of 8° from fixation. One of the four locations (randomly selected) was framed in dark gray (∼24 cd/m2), indicating the attention target (AT), i.e. the location which participants should aim to attend to endogenously. Note that no such attention target was indicated in the *Eye-only*, the *Hand-only*, and the *Eye-Hand* task. Once stable eye and finger fixation was detected within a 2.5° radius virtual circle centered on the fixation targets, four streams of 1/f spatial noise patches (radius 1.7°) appeared at the marked locations. Each noise stream consisted of randomly generated 1/f noise patches windowed by a symmetrical raised cosine (radius 1.7°, sigma 0.57), refreshing at 60 Hz (***Figure 1b***). After 400 −800 ms, two arrow cues appeared nearby the FT, indicating the eye and the hand movement targets (MT1 & MT2). The motor targets were selected randomly for each trial and could coincide with the attention target as well as with each other. The onset of the arrow cues was the go-signal for both movements, which had to be executed as fast and precise as possible. Participants reached towards either of the two potential motor targets while simultaneously making a saccade towards the other—at free choice. Note that in the *Attention-only* task no cues occurred, and in the single movement tasks (*Eye-only, Hand-only, Attention-Eye*, and *Attention-Hand*) only one arrow occurred— and only one movement was executed while the other effector remained at the fixation target. 100 −150 ms after cue onset (within the movement latency), one of the 1/f noise streams was briefly replaced by an orientation-filtered noise stimulus, showing a 40° clockwise or counterclockwise orientation. Participants were informed that this test signal would appear at the attention target location in 75 % of trials (in tasks without an attention target, the test was equally likely to appear at any of the four locations). After 50 ms the test was masked by the reappearance of non-oriented 1/f noise for another 700 ms (***Figure 1c*** provides an overview of stimulus timing). Afterwards, the screen turned blank and participants indicated via button press in a non-speeded manner whether they had perceived the orientation to be tilted clockwise or counterclockwise. They received auditory feedback for incorrect responses.

A threshold task preceded the experiment to ensure a consistent level of discrimination difficulty across participants. The threshold task visually matched the main experiment but no arrow cues were presented and participants were instructed to maintain eye and finger fixation. Furthermore, they were informed at which of the 4 locations the test would be presented in 100 % of trials. We used a procedure of constant stimuli and randomly selected the orientation filter strength *alpha* (corresponding to the visibility of the orientation tilt) out of six linear steps of filter widths. By fitting cumulative Gaussian functions to the discrimination performance via maximum likelihood estimation, we determined the filter width corresponding to 75% correct discrimination performance for each participant and used this value for the main experiment.

Participants performed 66 experimental blocks (2 *Attention-only*, 3 *Eye-only*, 2 *Hand-only*, 8 *Eye-Hand*, 11 *Attention-Eye*, 8 *Attention-Hand*, and 32 *Attention-Eye-Hand* blocks) of at least 66 trials each, resulting in a total of 4,356 trials per participant. We controlled online for violations of eye and finger fixation (outside 2.5° from FT before the cue onset), too short (<170 ms) or too long (>700 ms) movement latencies, and incorrect eye or hand movements (not landing within 2.5° from motor target center). Erroneous trials were repeated in random order at the end of each block. Overall, 567 ± 117 (mean ± SEM) trials per participant were repeated due to eye movement errors, 441 ± 73 due to finger movement errors.

Task, stimuli, and timing of **Experiment 2** were equivalent to Experiment 1, except that only one stream of 1/f noise patches was presented to which participants were endogenously attending throughout. The location of this noise stream (attention target; AT) again was indicated at the beginning of each block (either the upper right or the upper left location, randomly selected). As in the previous experiment, depending on the pre-block instruction, participants had to either exclusively attend to the noise stream (*Attention-only*), or attend to the noise stream and perform eye-(*Attention-Eye*), hand-(*Attention-Hand*), or simultaneous eye-hand-movements (*Attention-Eye-Hand*) to randomly selected motor target(s) indicated by centrally presented arrow cue(s). Unlike in Experiment 1, attention and motor targets never coincided. Furthermore, for each trial we randomly selected the orientation filter strength out of eight linear steps of filter widths (i.e. visibility level; *alpha* 5 to 75) and fitted cumulative Gaussian functions to the obtained group average discrimination performance via maximum likelihood estimation.

After an initial training (one block of 30 trials for each movement condition), participants performed 13 experimental blocks (3 *Attention-only*, 3 *Attention-Eye*, 3 *Attention-Hand*, and 4 *Attention-Eye-Hand* blocks) of at least 80 trials each, resulting in a total of 1,130 trials per participant. We controlled online for violations of eye and finger fixation (outside 2.5° from the FT before the cue onset), too short (<170 ms) or too long (>700 ms) movement latencies, and incorrect eye or hand movements (not landing within 2.5° from motor target center). Erroneous trials were repeated in random order at the end of each block. Overall, 145 ± 67 trials per participant were repeated due to eye movement errors, 130 ± 29 due to finger movement errors.

### Eye data pre-processing

We scanned the recorded eye-position data offline and detected saccades based on their velocity distribution [Engbert & Mergenthaler, 2006] using a moving average over twenty subsequent eye position samples. Saccade onset and offset were detected when the velocity exceeded or fell below the median of the moving average by 3 SDs for at least 20 ms. We included trials if a correct fixation was maintained within a 2.5° radius centered on FT until cue onset and landed within 2.5° from the cued location no later than 700 ms following cue onset, and if no blink occurred during the trial. In total we included 39,751 trials in the analysis of the behavioral results for Experiment 1 (on average 3,975 ± 79 trials per participant) and 6,015 trials (1,003 ± 29 per participant) for Experiment 2.

### Statistical analysis and data visualization

For Experiment 1, we determined percentage correct discrimination performance separately for each task and location, depending on the respective motor and attention target configuration. Whisker plots show single participant discrimination performance (represented by dots) averaged across participants (represented by black lines) and corresponding 95% confidence intervals (indicated by colored bars). All comparisons were contrasted to the average performance at the movement-irrelevant (non-target) locations in the respective task (referred to as “neutral” / “N”), unless otherwise stated. Psychometric functions for the four tasks of Experiment 2 were obtained by fitting cumulative Gaussian functions to the group average orientation discrimination performance via maximum likelihood estimation. For all statistical comparisons we used permutation tests to determine whether the performance between two conditions (e.g. at cued vs. uncued locations) differed significantly. We resampled our data to create a permutation distribution by randomly rearranging the labels of the respective conditions for each participant and computed the difference in sample means for 1000 permutation resamples (iterations). We then derived p-values by locating the actually observed difference (difference between the group-averages of the two conditions) on this permutation distribution, i.e. the p-value corresponds to the proportion of the difference in sample means that fell below or above the actually observed difference. Unless otherwise stated, all reported differences remained significant after Bonferroni multiple-comparison correction. All files will be available via the OSF database upon manuscript publication (https://osf.io/q8nbd).

## Acknowledgements

The authors are grateful to Andreas Oschwald and the members of the Deubel lab for helpful comments and discussions in the beer garden.

## Additional information

### Author contributions

Nina M. Hanning: Conceptualization, Methodology, Software, Validation, Formal Analysis, Investigation, Writing - Original Draft Preparation, Writing - Review & Editing, Visualization, Project Administration; Luca Wollenberg: Conceptualization, Investigation, Writing - Review & Editing; Donatas Jonikaitis: Conceptualization, Writing - Original Draft Preparation, Writing - Review & Editing; Heiner Deubel: Conceptualization, Writing - Review & Editing, Funding Acquisition, Supervision;

### Ethics

The protocols for the study were approved by the ethical review board of the Faculty of Psychology and Education of the Ludwig-Maximilians-Universität München (approval number 13_b_2015), in accordance with German regulations and the Declaration of Helsinki. All participants gave written informed consent.

## References

Anderson, B., Laurent, P., Yantis, S. (2011). Value-driven attentional capture. Proceedings of the National Academy of Sciences 108(25), 10367 – 10371.

Armstrong, K. M., & Moore, T. (2007). Rapid enhancement of visual cortical response discriminability by microstimulation of the frontal eye field. Proceedings of the National Academy of Sciences, 104(22), 9499–9504.

Aston-Jones, G. S., Desimone, R., Driver, J., Luck, S. J., & Posner, M. I. (1999). Attention. In “Fundamental Neuroscience”(MJ Zigmond, FE Bloom, SC Landis, JL Roberts, and LR Squire, eds.).

Baldauf, D., Cui, H., & Andersen, R. A. (2008). The posterior parietal cortex encodes in parallel both goals for double-reach sequences. Journal of Neuroscience, 28(40), 10081–10089.

Baldauf, D., Deubel, H. (2008). Visual attention during the preparation of bimanual movements. Vision Research, 48(4), 549 – 563.

Baldauf, D., & Deubel, H. (2009). Attentional selection of multiple goal positions before rapid hand movement sequences: An event-related potential study. Journal of Cognitive Neuroscience, 21(1), 18–29.

Baldauf, D., & Deubel, H. (2010). Attentional landscapes in reaching and grasping. Vision Research, 50(11), 999–1013.

Baldauf, D., Wolf, M., & Deubel, H. (2006). Deployment of visual attention before sequences of goal-directed hand movements. Vision Research, 46(26), 4355–4374.

Baruni, J., Lau, B., Salzman, C. (2015). Reward expectation differentially modulates attentional behavior and activity in visual area V4. Nature Neuroscience 18(11), 1656 – 1663.

Bichot, N., Rossi, A., Desimone, R. (2005). Parallel and serial neural mechanisms for visual search in macaque area V4. Science 308(5721), 529 – 534.

Bisley, J. W., & Goldberg, M. E. (2010). Attention, intention, and priority in the parietal lobe. Annual Review of Neuroscience, 33, 1–21.

Bisley, J. W., & Mirpour, K. (2019). The neural instantiation of a priority map. Current Opinion in Psychology, 29, 108–112.

Brainard, D. H. (1997). The psychophysics toolbox. Spatial Vision, 10(4), 433–436.

Bundesen, C. (1990). A theory of visual attention. Psychological Review, 97(4), 523–547.

Bundesen, C., Habekost, T., & Kyllingsbæk, S. (2011). A neural theory of visual attention and short-term memory (NTVA). Neuropsychologia, 49(6), 1446–1457.

Carrasco, M., Ling, S., & Read, S. (2004). Attention alters appearance. Nature neuroscience, 7(3), 308–313.

Carrasco, M. (2011). Visual attention: The past 25 years. Vision Research, 51(13), 1484–1525.

Ciaramitaro, V., Cameron, E., Glimcher, P. (2001). Stimulus probability directs spatial attention: an enhancement of sensitivity in humans and monkeys. Vision Research 41(1), 57 – 75.

Colby, C., Duhamel, J., Goldberg, M. (1996). Visual, presaccadic, and cognitive activation of single neurons in monkey lateral intraparietal area. Journal of Neurophysiology, 76(5), 2841 – 2852.

Cornelissen, F. W., Peters, E. M., & Palmer, J. (2002). The Eyelink Toolbox: eye tracking with MATLAB and the Psychophysics Toolbox. Behavior Research Methods, Instruments, & Computers, 34(4), 613–617.

Cosman, J., Lowe, K., Zinke, W., Woodman, G., Schall, J. (2018). Prefrontal control of visual distraction Current Biology 28(3), 414 - 420.e3.

Craighero, L., Fadiga, L., Rizzolatti, G., & Umiltà, C. (1999). Action for perception: a motor-visual attentional effect. Journal of Experimental Psychology: Human Perception and Performance, 25(6), 1673–1692.

Deubel, H. (2008). The time course of presaccadic attention shifts. Psychological Research, 72(6), 630–640.

Deubel, H., & Schneider, W. X. (1996). Saccade target selection and object recognition: Evidence for a common attentional mechanism. Vision Research, 36(12), 1827–1838.

Deubel, H., Schneider, W. X., & Paprotta, I. (1998). Selective dorsal and ventral processing: Evidence for a common attentional mechanism in reaching and perception. Visual Cognition, 5(1-2), 81–107.

Ekstrom, L. B., Roelfsema, P. R., Arsenault, J. T., Bonmassar, G., & Vanduffel, W. (2008). Bottom-up dependent gating of frontal signals in early visual cortex. Science, 321(5887), 414–417.

Engbert, R., & Mergenthaler, K. (2006). Microsaccades are triggered by low retinal image slip. Proceedings of the National Academy of Sciences, 103(18), 7192–7197.

Fecteau, J. H., & Munoz, D. P. (2006). Salience, relevance, and firing: a priority map for target selection. Trends in Cognitive Sciences, 10(8), 382–390.

Fernández A & Carrasco M (2020). Extinguishing exogenous attention via transcranial magnetic stimulation. Current Biology, 30, 1–7.

Found, A., Müller, H. (1996). Searching for unknown feature targets on more than one dimension: investigating a “dimension-weighting” account. Perception & Psychophysics 58(1), 88 – 101.

Gilster, R., Hesse, C., & Deubel, H. (2012). Contact points during multidigit grasping of geometric objects. Experimental Brain Research, 217(1), 137–151.

Godijn, R., & Theeuwes, J. (2003). Parallel allocation of attention prior to the execution of saccade sequences. Journal of Experimental Psychology: Human Perception and Performance, 29(5), 882 – 896.

Hanning, N. M., Aagten-Murphy, D., & Deubel, H. (2018). Independent selection of eye and hand targets suggests effector-specific attentional mechanisms. Scientific Reports, 8:9434.

Hanning, N. M., Deubel, H., & Szinte, M. (2019). Sensitivity measures of visuospatial attention. Journal of Vision, 19(12).

Hanning, N. M., Szinte, M., & Deubel, H. (2019). Visual attention is not limited to the oculomotor range. Proceedings of the National Academy of Sciences, 116(19), 9665–9670.

Hesse, C., & Deubel, H. (2011). Efficient grasping requires attentional resources. Vision Research, 51(11), 1223–1231.

Hesse, C., Schenk, T., & Deubel, H. (2012). Attention is needed for action control: further evidence from grasping. Vision Research, 71, 37–43.

Hoffman, J. E., & Subramaniam, B. (1995). The role of visual attention in saccadic eye movements. Perception & Psychophysics, 57(6), 787–795.

Ipata, A., Gee, A., Bisley, J., Goldberg, M. (2009). Neurons in the lateral intraparietal area create a priority map by the combination of disparate signals. Experimental Brain Research, 192(3), 479 – 488.

Itti, L., & Koch, C. (2001). Computational modelling of visual attention. Nature Reviews Neuroscience, 2(3), 194–203.

Jonikaitis, D., & Deubel, H. (2011). Independent allocation of attention to eye and hand targets in coordinated eye-hand movements. Psychological Science, 22(3), 339–347.

Jonikaitis, D., Klapetek, A., Deubel, H. (2017). Spatial attention during saccade decisions. Journal of neurophysiology 118(1), 149 – 160.

Jonikaitis, D., & Moore, T. (2019). The interdependence of attention, working memory and gaze control: behavior and neural circuitry. Current Opinion in Psychology, 29, 126–134.

Juan, C., Shorter-Jacobi, S., Schall, J. (2004). Dissociation of spatial attention and saccade preparation. Proceedings of the National Academy of Sciences of the United States of America, 101(43), 15541 – 15544.

Khan, A. Z., Song, J. H., & McPeek, R. M. (2011). The eye dominates in guiding attention during simultaneous eye and hand movements. Journal of Vision, 11(1).

Klein, R. M. (2000). Inhibition of return. Trends in Cognitive Sciences, 4(4), 138–147.

Kowler, E., Anderson, E., Dosher, B., & Blaser, E. (1995). The role of attention in the programming of saccades. Vision Research, 35(13), 1897–1916.

Kreyenmeier, P., Deubel, H., & Hanning, N. M. (2020). Theory of Visual Attention (TVA) in Action: Assessing Premotor Attention in Simultaneous Eye-Hand Movements. bioRxiv doi:10.1101/2020.01.08.898932

Land, M., Lee, D. (1994). Where we look when we steer. Nature 369(6483), 742 – 744.

Land, M., McLeod, P. (2000). From eye movements to actions: how batsmen hit the ball. Nature Neuroscience 3(12), 1340 – 1345.

Lovejoy, L. P., & Krauzlis, R. J. (2010). Inactivation of primate superior colliculus impairs covert selection of signals for perceptual judgments. Nature Neuroscience, 13(2), 261–266.

Mazer, J., Gallant, J. (2003). Goal-related activity in V4 during free viewing visual search: evidence for a ventral stream visual salience map. Neuron, 40(6), 1241 – 1250.

McAdams, C., Maunsell, J. (1999). Effects of attention on orientation-tuning functions of single neurons in macaque cortical area V4. The Journal of Neuroscience 19(1), 431 – 441.

McPeek, R., Skavenski, A., Nakayama, K. (2000). Concurrent processing of saccades in visual search. Vision Research 40(18), 2499 – 2516.

Montagna, B., Pestilli, F., & Carrasco, M. (2009). Attention trades off spatial acuity. Vision Research, 49(7), 735–745.

Montagnini, A., & Castet, E. (2007). Spatiotemporal dynamics of visual attention during saccade preparation: Independence and coupling between attention and movement planning. Journal of Vision, 7(14).

Morawetz, C., Holz, P., Baudewig, J., Treue, S., Dechent, P. (2007). Split of attentional resources in human visual cortex. Visual Neuroscience 24(6), 817 – 826.

Moore, T., & Armstrong, K. M. (2003). Selective gating of visual signals by microstimulation of frontal cortex. Nature, 421(6921), 370–373.

Moore, T., & Fallah, M. (2004). Microstimulation of the frontal eye field and its effects on covert spatial attention. Journal of Neurophysiology, 91(1), 152–162.

Moran, J., & Desimone, R. (1985). Selective attention gates visual processing in the extrastriate cortex. Science, 229(4715), 782–784.

Müller, H. J., & Krummenacher, J. (2006). Visual search and selective attention. Visual Cognition, 14(4-8), 389–410.

Müller, H. J., & Rabbitt, P. M. (1989). Reflexive and voluntary orienting of visual attention: time course of activation and resistance to interruption. Journal of Experimental Psychology: Human Perception and Performance, 15(2), 315.

Munoz, D., Everling, S. (2004). Look away: the anti-saccade task and the voluntary control of eye movement. Nature Reviews Neuroscience 5(3), 218 – 228.

Niebergall, R., Khayat, P., Treue, S., Martinez-Trujillo, J. (2011). Multifocal attention filters targets from distracters within and beyond primate MT neurons’ receptive field boundaries. Neuron 72(6), 1067 – 1079.

Pelli, D. G. (1997). The VideoToolbox software for visual psychophysics: Transforming numbers into movies. Spatial Vision, 10(4), 437–442.

Peters, R., Iyer, A., Itti, L., Koch, C. (2005). Components of bottom-up gaze allocation in natural images. Vision Research 45(18), 2397 – 2416.

Posner, M. I. (1980). Orienting of attention. Quarterly Journal of Experimental Psychology, 32(1), 3–25.

Reynolds, J. H., & Chelazzi, L. (2004). Attentional modulation of visual processing. Annu. Rev. Neurosci., 27, 611–647.

Rizzolatti, G., Riggio, L., Dascola, I., & Umiltá, C. (1987). Reorienting attention across the horizontal and vertical meridians: evidence in favor of a premotor theory of attention. Neuropsychologia, 25(1), 31–40.

Rolfs, M., Lawrence, B. M., & Carrasco, M. (2013). Reach preparation enhances visual performance and appearance. Philosophical Transactions of the Royal Society B: Biological Sciences, 368(1628), 20130057.

Rolfs, M., & Szinte, M. (2016). Remapping attention pointers: linking physiology and behavior. Trends in Cognitive Sciences, 20(6), 399–401.

Saber, G., Pestilli, F., Curtis, C. (2015). Saccade planning evokes topographically specific activity in the dorsal and ventral streams. Journal of Neuroscience 35(1), 245 – 252.

Schafer, R., Moore, T. (2007). Attention governs action in the primate frontal eye field. Neuron, 56(3), 541 – 551.

Schneider, W. X. (1995). VAM: A neuro-cognitive model for visual attention control of segmentation, object recognition, and space-based motor action. Visual Cognition, 2(2-3), 331–376.

Schütz, A. C., Trommershäuser, J., & Gegenfurtner, K. R. (2012). Dynamic integration of information about salience and value for saccadic eye movements. Proceedings of the National Academy of Sciences, 109(19), 7547–7552.

Serences, J. T. (2008). Value-based modulations in human visual cortex. Neuron, 60(6), 1169–1181.

Steinmetz, N., Moore, T. (2014). Eye movement preparation modulates neuronal responses in area V4 when dissociated from attentional demands. Neuron 83(2), 496 – 506

Theeuwes, J., & Burger, R. (1998). Attentional control during visual search: the effect of irrelevant singletons. Journal of Experimental Psychology: Human Perception and Performance, 24(5), 1342–1353.

Thompson, K. G., & Bichot, N. P. (2005). A visual salience map in the primate frontal eye field. Progress in Brain Research, 147, 249–262.

Theeuwes, J., Kramer, A., Hahn, S., Irwin, D. (1998). Our eyes do not always go where we want them to go Psychological science 5(5), 379 – 385.

Treisman, A. (1982). Perceptual grouping and attention in visual search for features and for objects. Journal of Experimental Psychology: Human Perception and Performance, 8(2), 194–214.

